# Slow change blindness from serial dependence

**DOI:** 10.1101/2025.10.02.679619

**Authors:** Haley G. Frey, David Whitney

## Abstract

Slow change blindness, when attentive observers fail to notice large changes that happen gradually, raises questions about how visual information is combined across time. One plausible integration strategy is serial dependence: blending information from the recent past into current perception. Here, we investigate serial dependencies in perception of a cartoon object that slowly changes hue. In a one-shot experiment, observers each viewed a single trial with a random degree of hue change and provided one hue judgement response. Across participants, the entire morph was probed. Observers’ hue reports revealed an overall bias towards the past that increased in magnitude as more of the morph was experienced. In three follow-up experiments, we verified that observers experienced slow change blindness, confirmed that the bias was serial dependence, and replicated the results with a repeated-trials design. Overall, we provide evidence that serial dependence actively biases perception during gradual changes, producing slow change blindness.

**Teaser:** By strategically integrating recent and current visual information, observers experience a slowly changing stimulus as stable.

## Introduction

Attentive observers fail to notice otherwise obvious changes when the change takes place gradually, a phenomenon called slow change blindness (Simons et al., 2000). Such changes include the appearance or disappearance of an object (Simons et al., 2000), rotation in scene orientation (Hollingworth & Henderson, 2004), and drift in the color or hue (David et al., 2006; Frey, Koenig, He, et al., 2024; Simons et al., 2000). Slow change blindness persists even when the change region is large, occurs in full view, and is centrally located (Frey, Koenig, He, et al., 2024) and surprisingly, even when the change involves highly relevant features such as facial expression (David et al., 2006) or age (Manassi & Whitney, 2022).

This striking inability to notice large changes raises questions about how perception is generated across time. Past studies have offered explanations of slow change blindness, often borrowed from classic change blindness paradigms (e.g., flicker paradigm, Rensink et al., 1997; mud-splash paradigm, O’Regan et al., 1999). Most of these hypotheses revolve around the notion that there are distinct pre- and post-change stimulus states and that change blindness results from a) a failure to encode one or both states (implicit updating hypothesis, Hollingworth & Henderson, 2004; world as external memory, O’Regan, 1992) or b) a failure to compare the contents of the two encoded states (Mitroff, 2004; Smith, Lamont, & Henderson, 2012). These proposals are sometimes unsatisfying: comparing two scenes or objects is a fundamentally different task for the visual system than continuously representing objects or scenes during which there are no clear pre- and post-change states. Classic change perception is a task that relies on attention and memory, while slow change perception is a task that involves continuous and strategic integration of information over time. Therefore, while classic change blindness can result from misplaced attention or insufficient memory representations, slow change blindness may unfold because of the nature of the integration mechanisms involved in online perception.

One integration method that would plausibly lead to slow change blindness is a continuous reliance on the recent past – a kind of “active perceptual serial dependence” (Manassi & Whitney, 2022). Serial dependence is a phenomenon in which observers’ perception of a current stimulus is biased towards recently seen stimuli, and is strongest when successive stimuli are more similar to one another (Cicchini et al., 2024; Kiyonaga et al., 2017a; Manassi et al., 2023; Pascucci et al., 2023). It is characterized by a spatiotemporal integration operator, the Continuity Field (Manassi & Whitney, 2024), which is agnostic about the level of implementation and could lead to serial dependencies at any or all stages of visual processing. Serial dependence is consistent with predictive coding and Bayesian optimal theories of visual processing (Cicchini et al., 2024; Kersten et al., 2004), yet could also be heuristic or a result of an interplay between encoding and decoding strategies (Cicchini et al., 2024; Teng & Kravitz, 2019; Zhang & Lewis-Peacock, 2024). Given the correlation of natural world statistics across time (Dong & Atick, 1995), assuming continuity and basing current perception on the recent past is a computationally efficient strategy that can improve vision during uncertainty (Cicchini et al., 2018). For example, serially dependent perception can promote object constancy throughout changing viewing angles, lighting conditions, and head and object positions (Biederman, 1987; DiCarlo & Cox, 2007; Palmeri & Gauthier, 2004). Although the evidence for serial dependence comes from trends across several distinct trials, it would make sense for the visual system to employ a similar strategy during continuous viewing. Given that serial dependence in perception is modulated by both similarity and time, it stands to reason that a series of slight adjustments to an object across time should lead to a series of serially dependent percepts. Across time, these continuously biased percepts could camouflage a slow change (Manassi & Whitney, 2022), leading to slow change blindness.

Of the handful of studies which have investigated slow change blindness, three of them provide results that could be consistent with serial dependence. After viewing an image where a face slowly changes expression, observers (unaware of the change) report a facial expression that reflects the face as it had been about three seconds earlier in the morph rather than the actual facial expression that they had just seen (Laloyaux et al., 2008). For example, if the face morphed from 100% neutral/0% happy to 0% neutral/100% happy, the observer was most likely to report the 25% neutral/75% happy face as the face they had seen (Laloyaux et al., 2008). In another study, observers who viewed a short morph reported the age of a face as much younger than truth when the preceding face morphed from young to old and reported the age as much older than truth when the preceding face morphed from old to young (Manassi &

Whitney, 2022), indicating that they were unaware of the extent of the age change. This study did not explicitly ask observers if they noticed that the face had changed age. The authors replicated this trend using morphs between male and female faces, showing that observers’ reports depended on the preceding morph (Manassi & Whitney, 2022). A more recent study found that observers (once again, blind to the change) who viewed a color-changing object, tended to report that the color presented midway through the morph best matched what they had just seen (Frey, Koenig, Block, et al., 2024). In each of the three studies described, observers’ reports of the final stimulus state were biased towards the past and depended on the particular preceding stimulus states.

Though the results of these previous studies could be consistent with a serially dependent integration process, these studies cannot make claims about moment-to-moment perception of a continuously changing object, nor can they rule out other potential integration methods. To disentangle moment-to-moment perception from morph-long reflective percepts or memories, a handful of other possibilities must be considered. First, it is possible that observers simply demonstrate hysteresis: that their perception updates continuously and is unbiased but lags the presented stimulus state. An observer reporting their perception after viewing a slowly changing morph may report an earlier state simply because their perception has not yet been updated to reflect the most recent stimulus, not because their perception includes information from the past. Second, it is possible that without any awareness of the change, observers calculate and report the average stimulus state when they are prompted to make a judgement about what they have just seen. Temporal ensemble perception could be an example of this (Haberman et al., 2009; Khayat et al., 2023; Whitney & Leib, 2018). Manassi & Whitney (2022) provide evidence against this by showing a change in the degree of perceptual bias when the task-irrelevant preceding morph traverses the same age range via discrete steps instead of continuously, but these videos no longer present the same task to the visual system (the steps may trigger a resetting of reference frame or draw attention to the change).

Importantly, the studies discussed above only probe perception of a static stimulus once after observers have experienced the full task-irrelevant morph; they do not measure instantaneous perception of a continuously changing stimulus. This leaves some crucial questions still unanswered: what is the perception of a slowly changing object at each moment in time and how is it generated?

Here, we investigate perception at each moment of a slow change, rule out hysteresis and averaging, and provide support for a role of active perceptual serial dependence in the phenomenon of slow change blindness. In the following studies, we measure perception at each moment in a morph by accumulating one-shot hue reports from a large number of observers who each viewed a different extent of the total slow hue change. To preview our results, we find that overall reported hue perception is biased towards the past (Experiments 1 and 2).

Additionally, and more importantly, we find that the bias increases as more of the morph is experienced, consistent with a dependence on the preceding information at each moment (Experiments 1 and 2). By explicitly asking whether observers noticed the change, we verify that the resulting illusion of stability (Manassi & Whitney, 2022) produces slow change blindness (Experiment 2). We show that reducing serial dependence via a change in object identity reduces the bias in hue reports, confirming that the measured bias is indeed serial dependence (Experiment 3). Lastly, we show that observers who experience repeated trials report a smaller yet still significant bias, suggesting that the serial dependence persists despite knowledge of the task (Experiment 4). In sum, these experiments overcome the limitations of previous work to yield results which strongly support the idea that active serial dependence underlies slow change blindness.

## Experiment 1

### Method

#### Participants

201 participants were recruited through Prolific (Palan & Schitter, 2018, www.prolific.com) and completed the study online through Pavlovia (Peirce et al., 2019; www.pavlovia.com) for pay at the rate of $10/hr. Participants were required to use a desktop computer but there was no other exclusion criteria. The study was approved by the University of California, Berkeley IRB and informed consent was given by each participant before beginning the study.

#### Procedure

The goal of this experiment was to obtain a continuous measure of perception to a slowly changing object in a scene. To eliminate any repeated measure concerns and biases, this “continuous” measurement was achieved by stitching together single participant one-shot responses to various degrees of change. The slow change in this experiment involved a color change of a large, centrally located cartoon couch in a living room scene. Cartoon images were chosen due to their ease in editing, plausible range of colors, and scene-like structure which introduced meaning to the color-changing object. Specifically, hue was changed here because it could be easily manipulated and quantified; each intermediate stage throughout the morph was a valid state (as opposed to a semi-transparent object that would exist during a slow appearance or disappearance) and is a salient feature that is likely to have been in the observers’ awareness despite blindness to the change. Throughout the rest of the manuscript, this slow color change will be referred to as a hue change because this was the aspect of color that was changed.

Each participant experienced a single trial where they viewed the colored couch (“test couch”) for a random duration, dictated by the randomly assigned degree of hue change.

Participants reported the hue of the couch immediately after the morph ended – making a single hue response (Figure 1a). Participants were unaware that the couch could change hue. Before beginning, participants were given the following instructions: “You’ll see a colored image of a couch in a living room for a random duration, up to 40 seconds. After that, you’ll be asked to report the color of the couch. Try to be as accurate as possible! When you are ready, press ‘space’.” Therefore, though observers did not know that the couch could change in hue, they did know that they would be asked to report on the color of the couch they viewed. After the couch image disappeared from the screen, a color wheel and corresponding “response couch” (the same gray as the monitor background) were presented. Participants were given the following instructions: “What color was the couch? Hover over the color wheel to adjust the color of the couch. When you have reached your choice, click with the mouse.” As the participants hovered over the wheel, the center of the color wheel and the “response couch” were updated to reflect the current color under their mouse. This feature was included to account for any simultaneous-color-contrast effects resulting from the other colors present in the scene and to ensure as accurate a response as possible. The color wheel was defined in HSV space and contained all hues (0-359 degrees) at a saturation of .40 and value of 0.82. The color wheel contained more colors than the participant was exposed to during the trial, but all colors that the participant saw were contained in the color wheel. The orientation of the color wheel (which color was at the top) was randomized for each participant. Participants selected their choice by clicking with the mouse. Morph duration (time), morph distance (hue), actual ending hue, and selected ending hue were recorded for each participant’s trial.

**Figure 1.**
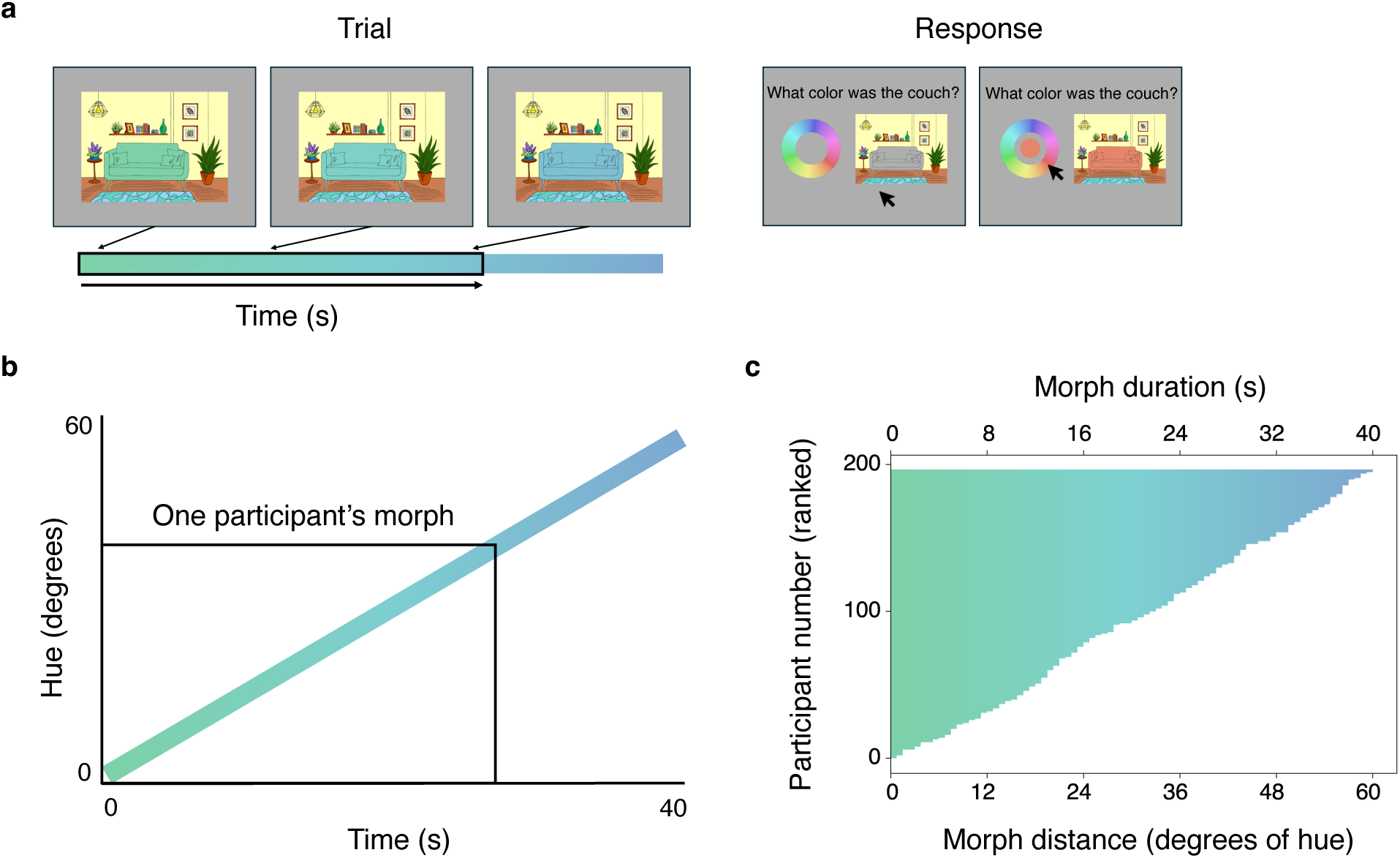
Experiment 1 stimulus & design. **a.** The trial consisted of a test couch that changed hue across time. On the left, the test couch is shown in its initial state. The color gradient below the couch represents the maximum hue change possible, with the black outline showing the random subset of the morph that this example participant experienced. The arrows point to the current hue of the couch in each frame. In this example, the rightmost couch is the last one shown before the participant is asked to make a color judgement. On the right is the response screen. Participants used their mouse to hover over the color wheel. As they hovered over the wheel (right), the center of the color wheel and the response couch matched the color under the mouse. Participants clicked with their mouse to select their choice. **b.** A plot showing stimulus motion (hue change) across time, with an example participant’s morph outlined in black. **c.** The extent of the morph experienced by each participant. The bottom x-axis shows the morph distance in terms of hue, with the corresponding duration in time on the top x-axis. The y-axis shows individual participants, ranked in terms of exposure duration. A single row represents the hue change that a single observer saw, with the rightmost edge of the bar indicating the actual ending hue at the moment the observer was asked to make a response.

In this experiment, the starting couch hue was always 150 degrees (in HSV space), and the ending hue was randomly assigned up to a maximum ending hue of 210 degrees. The hue always changed at a rate of 1.5 degrees/s (Figure 1b). Therefore, each participant experienced a total hue change between 0.75 and 60 degrees, and, as a result, a different video duration (s), depending on the amount of hue change. Figure 1c shows that across participants, the entire range of hue distances relative to the starting point were covered. Due to the differences in the degree of hue change - and therefore stimulus duration - across participants, each participant experienced a different amount of preceding hue information prior to entering their color report. To reiterate, each individual participant was exposed to a different and random degree of hue change during the single trial (Figure 1c). A strength of this one-shot, single trial approach is that participants were completely naïve; they had no knowledge that the hue could change, no reference of the color range beforehand, and limited knowledge of their reporting task.

#### Preprocessing

For each participant, error was calculated as selected ending hue – actual ending hue. Raw errors spanned the range −195 to 123 but were converted to fit in the range of −180 to +180. Errors less than −180 are equivalent to 360 + error and errors greater than 180 are equivalent to error – 360. Participants with an error greater than three times the standard deviation above the mean or less than three times the standard deviation below the mean were excluded from analysis and are not depicted in any figures (4 participants; inclusion or exclusion of these participants does not meaningfully change the results).

Before beginning the task, participants were asked to report whether they had normal or corrected-to-normal (wearing glasses or contacts) vision, and whether they were colorblind. From these responses, participants were split into an “impaired” (N = 39) and a “sighted” (N = 158) group. A permutation analysis was conducted to determine whether the difference in mean error between the “impaired” and “sighted” groups was significant. The empirical mean difference (−0.148) was compared to a permuted null distribution, testing the null hypothesis that there is no difference between the means. To create the null distribution, the experiment labels were randomly permuted 10,000 times and the slope was recalculated at each permutation. The resulting p-value was .519 (α = 0.0167), indicating that the empirical mean difference was not significantly different than zero. Therefore, no data was excluded based on sight during the following analyses.

## Results

Figure 2a shows selected hue as a function of relative hue distance from start. A simple linear regression was fit to the empirical data: selected = 145.9 + 0.684*distance (solid black). The model verified that as hue distance travelled increased, selected hue increased as well, r(195) = .506, t = 8.19, p < .001. The dotted black line is a unity line representing ground truth: the actual hue at each distance. To determine whether the slope of the empirical data differed from the ground truth (solid vs dotted), an interaction analysis was conducted. A multiple linear regression model (selected ∼ distance*group) was fit to the combination of the ground truth (group 0) and empirical (group 1) data. The model yielded a significant interaction term (distance:group), confirming that the effect of distance on selected hue differed depending on the group, t = −3.78, p < .001. Specifically, the slope of the selected hue data was significantly flatter than the slope of the actual hue data. The model also yielded a significant main effect of distance (t = 16.9 p < .001) and a non-significant main effect of group (t = −1.36, p = .174).

**Figure 2.**
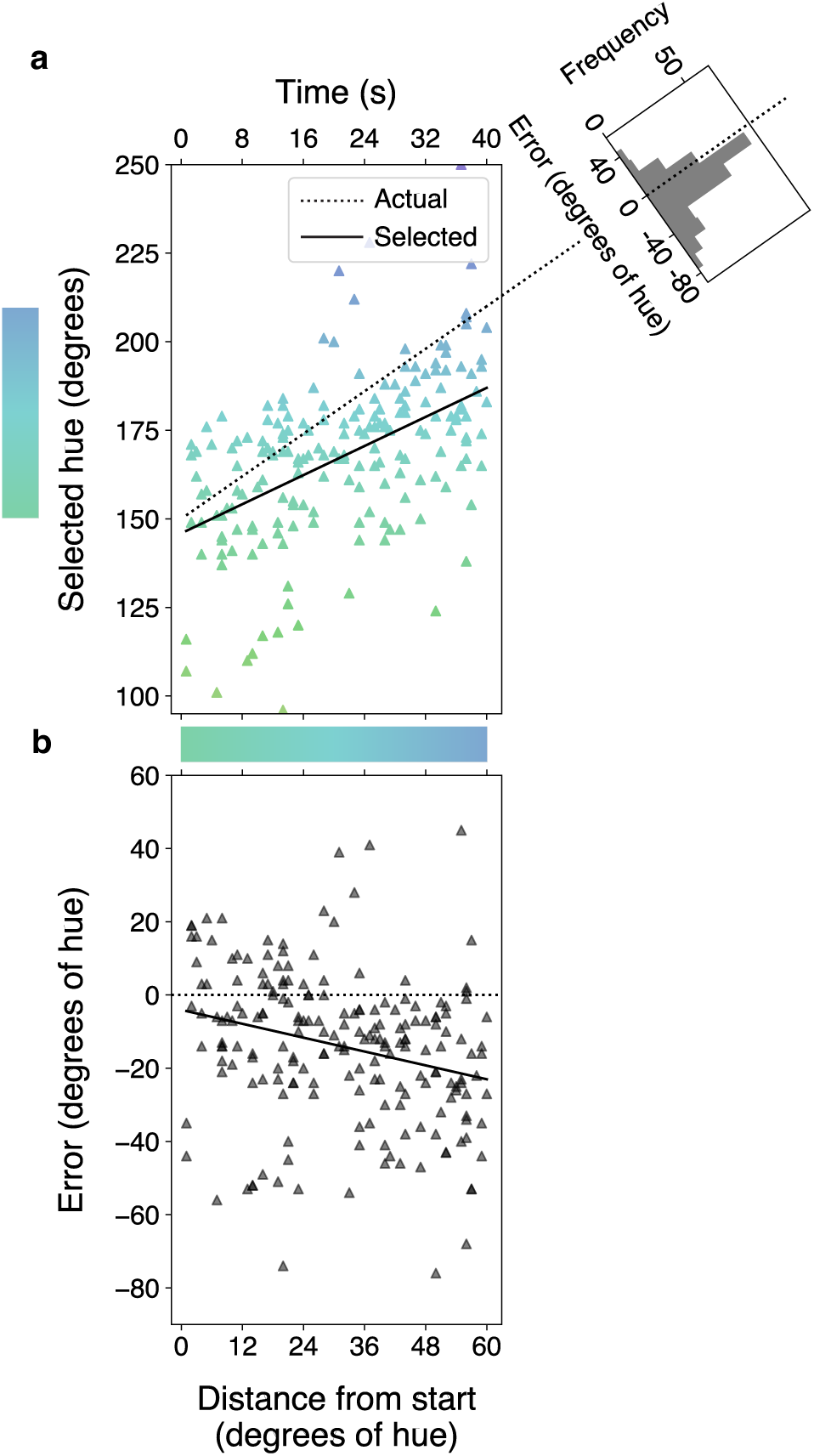
Experiment 1 results. **a.** Selected hue is plotted as a function of the hue distance travelled. Equivalent time is shown on the top x-axis. One triangle is one participant. The x-axis position reflects the relative morph endpoint, and the horizontal gradient bar shows the actual color at each hue distance. The y-axis position, also the color of the triangle, is the selected hue. The dotted black line is a unity line representing ground truth: the actual hue at each distance. Any triangle that does not fall on the dotted line indicates a nonzero error. The solid black line is a regression line fit to the selected hue data. The histogram in the top right corner plots the distribution of errors (selected hue – actual hue). The extended unity line represents zero error. **b.** Error is plotted as a function of hue distance from start, with equivalent time on the top x-axis. Each triangle represents one participant. The dotted black line is ground truth and the solid black line is a linear regression fit to the error data.

A histogram of errors collapsed across the morph, mean of −14.0 (SD = 20.6), is shown in the top right of Figure 2a. A one-sample t test determined that the mean error was significantly less than zero, t(196) = −9.56, p < .001. Further, a chi-square goodness-of-fit test confirms that the distribution of errors is significantly different than chance (equal frequency of responses above and below zero), 𝜒^2^ (1, 𝑁 = 193) = 63.8, 𝑝 < .001: the majority of errors are negative.

Figure 2b shows error as a function of hue distance from start. There was a negative relationship between hue distance and error (error = −4.08 + −0.316*distance): as the endpoint of the morph increased, error became more negative. A linear regression model revealed this trend was significant, r(195) = −.261, t = −3.78, p < .001.

## Discussion

Observers’ selected hues were predicted by the hue distance travelled (analogous to time; Figure 2a), indicating that observers update their representation across time, and do not simply “stick” with the initial stimulus state (as might be expected if there was anchoring, or if one used the world as external memory, O’Regan, 1992). The significance of this relationship demonstrates that observers are not choosing hues at random: their choices are related to the morph they experienced. However, the empirical slope (0.684) was significantly flatter than the ground truth slope (1), meaning that observers’ reports of the ending hue were 68.4% of the actual ending hue. In terms of time, the selected hue at time X matched the actual hue from an earlier timepoint. Despite updating their representations, observers’ representations are nonetheless clearly biased towards the past. The nonuniform distribution of errors and the negative mean error emphasized the tendency of participants to report an ending hue that is “less than” the true ending hue.

The pattern of the selected hues across the morph supports serial dependence but, examining error as a function of morph distance rules out alternative explanations more intuitively (Figure 2b). Both hysteresis (a stable lag in observers’ reports) and the implicit updating hypothesis (Hollingworth & Henderson, 2004) predict a constant error as the morph progresses. Though the predicted magnitude of the error differs (hysteresis tolerates a larger negative error while the implicit updating hypothesis expects a negative error closer to zero), both explanations predict a near-zero slope. However, error became significantly more negative as the hue distance traveled increased (slope = −0.408; Figure 2b). As the actual hue became more different than the starting hue, observers became less accurate in their reports. This non-zero slope rules out hysteresis and implicit updating and provides strong evidence for serial dependence. We conclude that observers continuously integrate information from the recent past, leading to a dynamic percept that becomes increasingly different from reality as the stimulus continues to change.

In Experiment 1, we replicated previous work demonstrating that after viewing a slowly changing stimulus, observers report a previous stimulus state as being the current ending state (Frey, Koenig, Block, et al., 2024; Laloyaux et al., 2008; Manassi & Whitney, 2022). Importantly, by stitching together one-shot responses to different endpoints along a single morph, we revealed a pattern of perceptual reports across time that allowed a more in depth analysis of perception over time. We found that the magnitude of the bias in perception (error) depends on the preceding information, with a longer morph exposure producing a greater bias. These results provide evidence in favor of serial dependence and rule out hysteresis and implicit updating as alternative explanations.

## Experiment 2

Though previous studies reported extremely high rates of slow change blindness to similar slow color-changing stimuli (Frey, Koenig, Block, et al., 2024; Frey, Koenig, He, et al., 2024), we could not be certain that our observers were blind to the change in Experiment 1. In Experiment 2, we repeated the above procedure and added a question asking explicitly whether the observer noticed a change in the couch color. With a measure of slow change blindness rate, we could confirm that these biases are present when observers experience slow change blindness. Further, we would be able to investigate whether there is a relationship between the reported bias and whether an observer reported noticing the change. We used this replication as an opportunity to test a new hue range, allowing us to verify that our initial results were not an artifact of the particular hues used. Finally, to alleviate any concern that observers merely reported the average hue perceived during the morph, we updated the wording of our test question to ask more specifically about the final state of the couch.

## Method

### Participants

201 new participants were recruited through Prolific (Palan & Schitter, 2018; www.prolific.com) and completed the study online through Pavlovia (Peirce et al., 2019; www.pavlovia.com) for pay at the rate of $10/hr. Participants were required to use a desktop computer but there was no other exclusion criteria. The study was approved by the University of California, Berkeley IRB and informed consent was given by each participant before beginning the study.

### Procedure

The procedure was identical to Experiment 1 except as described below. The instructions were altered slightly to better align with the updated test question. The new instructions read: “You’ll see a colored image of a couch in a living room for a random duration, up to 40 seconds. Please pay attention to the couch. At some point a black outline will surround the image and you will be asked a question about the couch. Try to be as accurate as possible! When you are ready, press ‘space’.” As in Experiment 1, the “test couch” was shown on the screen for a random duration, dictated by the randomly assigned degree of hue change. For the last second of the morph (1.5 degrees of hue change) and two additional seconds of static hue (the ending hue), a black outline box and the words “What is the color of THIS couch?” appeared on the screen (Figure 3a). This question was intended to prompt participants to specifically report the color of the couch in front of them rather than reporting an average from across the morph or any earlier state, reducing the contributions of memory. The “test couch” and question then disappeared, and the color wheel, “response couch”, and text were presented to collect the color judgement. After the participant submitted their color choice, they were then asked:

**Figure 3.**
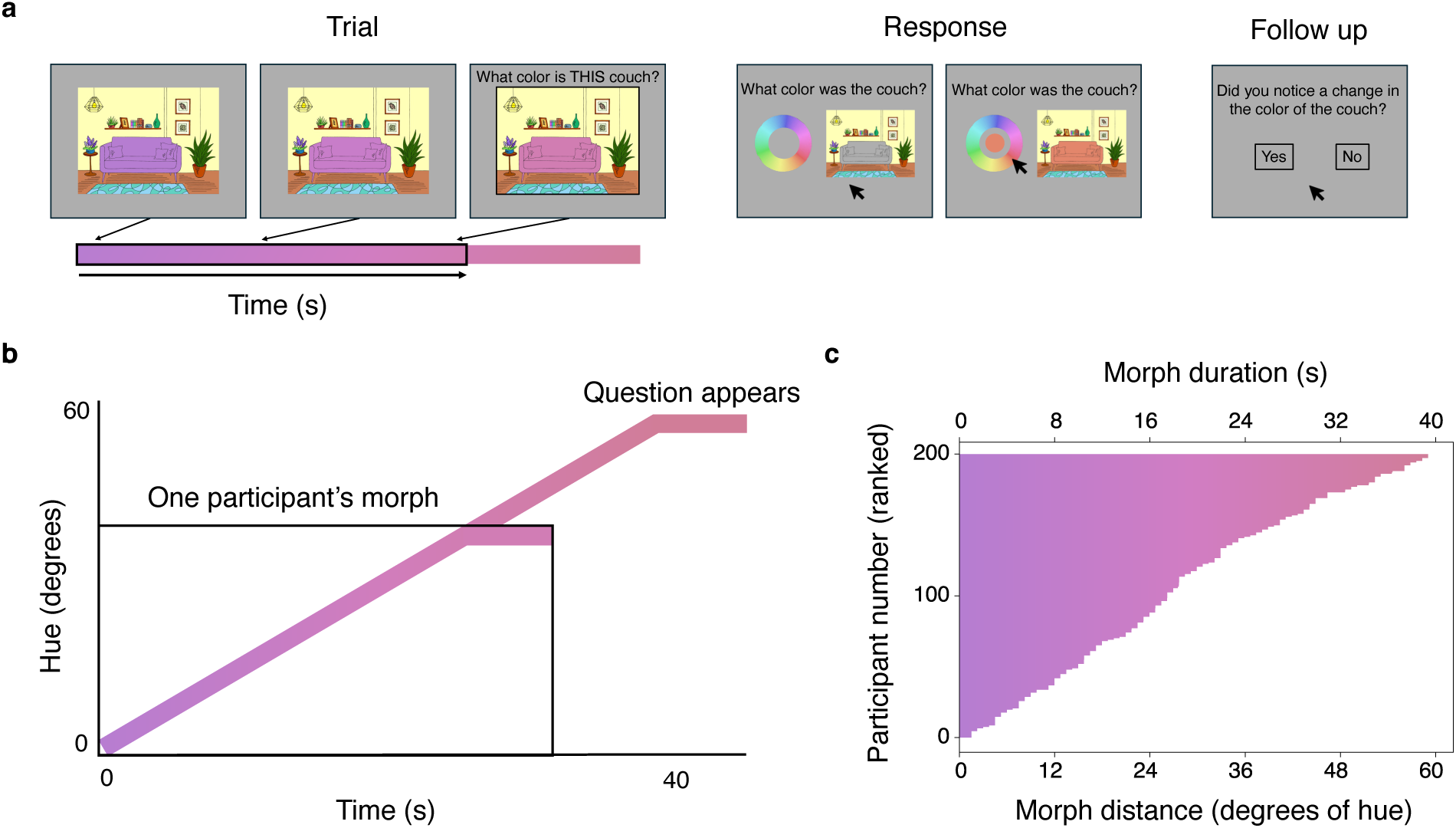
**Experiment 2 stimulus & design**. **a.** The trial consisted of a test couch that changed hue across time. On the left, the test couch is shown in its initial state. The color gradient below the couch represents the maximum hue change possible, with the black outline showing the random subset of the morph that this example participant experienced. The arrows point to the current hue of the couch in each frame. The rightmost frame shows that the question appeared with the couch still displayed for the last second of the morph and two additional seconds. On the right is the response screen. Participants used their mouse to hover over the color wheel. As they hovered over the wheel (right), the center of the color wheel and the response couch matched the color under the mouse. Participants clicked with their mouse to select their choice. After making their choice, participants were asked whether they noticed a change during the trial. **b.** A plot showing stimulus motion (hue change) across time, with an example participant’s morph outlined in black. **c.** The extent of the morph experienced by each participant. The bottom x-axis shows the morph distance in terms of hue, with the corresponding duration in time on the top x-axis. The y-axis shows individual participants, ranked in terms of exposure duration. A single row represents the hue change that a single observer saw, with the rightmost edge of the bar indicating the actual ending hue at the moment the observer was asked to make a response.

“While viewing the couch on the screen, did you notice any change in the color of the couch?” and responded by clicking a “yes” box or a “no” box on the screen. Morph duration (time), morph distance (hue), actual ending hue, selected ending hue, and whether the change was noticed were recorded for each participant’s trial.

In this iteration, the starting couch hue was always 280 degrees (in HSV space), and the ending hue was randomly assigned up to a maximum ending hue of 340 degrees. The maximum hue distance was 60 degrees and the rate of change was constant across participants at 1.5 degrees/second (Figure 3b). Each individual participant was exposed to a different and random degree of hue change during the single trial (Figure 3c).

### Preprocessing

For each participant, error was calculated as reported hue – actual hue. Raw errors spanned the range −120 to 70. Participants with an error greater than three times the standard deviation above the mean or less than three times the standard deviation below the mean were excluded from analysis and are not depicted in any figures (2 participants).

Before beginning the task, participants were asked to report whether they had normal or corrected-to-normal (wearing glasses or contacts) vision, and whether they were colorblind. From these responses, participants were split into an “impaired” (N = 55) and a “sighted” (N = 144) group. A permutation analysis was conducted to determine whether the difference in mean error between the “impaired” and “sighted” groups was significant. The empirical mean difference (1.22) was compared to a permuted null distribution, testing the null hypothesis that there is no difference between the means. To create the null distribution, the experiment labels were randomly permuted 10,000 times and the slope was recalculated at each permutation.

The resulting p-value was .354 (α = 0.0167), indicating that the empirical mean difference was not significantly different than zero. Therefore, no data was excluded based on sight during the following analyses.

## Results

Figure 4a shows selected hue as a function of relative hue distance from start. The solid black line, a linear regression fit to the empirical data (selected = 280.0 + 0.353*distance), revealed that morph distance was a significant predictor of selected hue, r(197) = .265, t = 3.84, p < .001. The dotted black line is a unity line representing ground truth. An interaction analysis was conducted to determine whether the slope of the empirical data differed from the ground truth (solid vs dotted). A multiple linear regression model (selected ∼ distance*group) was fit to the combination of the ground truth (group 0) and empirical (group 1) data. The model yielded a significant interaction term (distance:group), confirming that the effect of distance on selected hue differed depending on the group, t = −7.04, p < .001. Specifically, the slope of the selected hue data was significantly flatter than the slope of the actual hue data. The model also yielded a significant main effect of distance (t = 15.4, p < .001) and a non-significant main effect of group (t = 0.008, p = .993).

**Figure 4.**
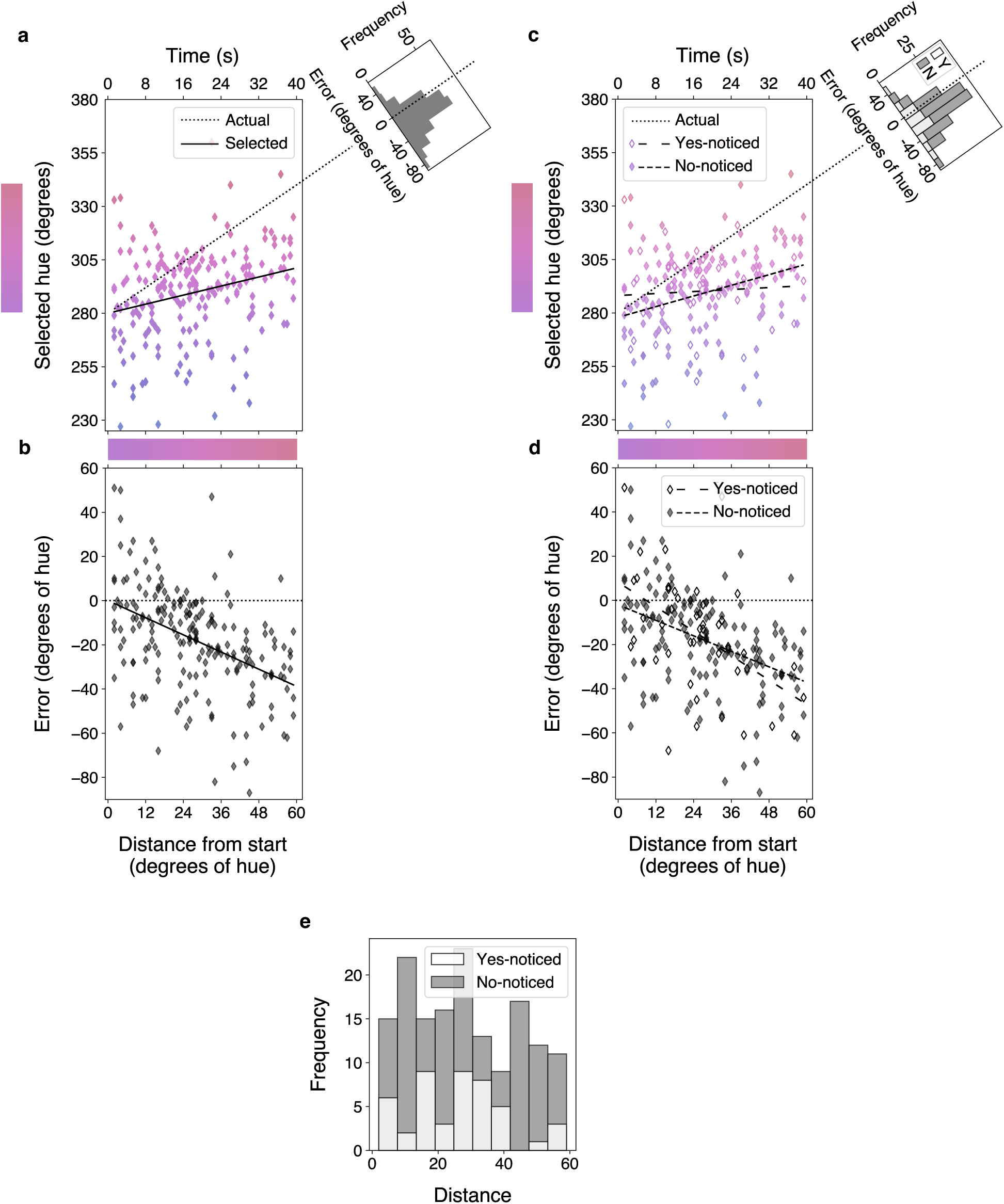
Experiment 2 results. **a.** Selected hue is plotted as a function of the hue distance travelled. Equivalent time is shown on the top x-axis. One diamond is one participant. The x-axis position reflects the relative morph endpoint, and the horizontal gradient bar shows the actual color at each hue distance. The y-axis position, also the color of the diamond, is the selected hue. The dotted black line is a unity line representing ground truth: the actual hue at each distance. Any diamond that does not fall on the dotted line indicates a nonzero error. The solid black line is a regression line fit to the selected hue data. The histogram in the top right corner plots the distribution of errors (selected hue – actual hue). The extended unity line represents zero error. **b.** Error is plotted as a function of hue distance from start, with equivalent time on the top x-axis. Each diamond represents one participant. The dotted black line is ground truth and the solid black line is a linear regression fit to the error data. **c & d.** Replicate panels a & b with data split by yes-noticed (open; long dash) and no-noticed (filled; short dash). **e.** Frequency of noticed-response (yes or no) as a function of morph distance.

A histogram of errors collapsed across the morph, mean of −17.8 (SD = 22.9), is presented in the top right of Figure 4a. A one-sample t test confirmed that the mean was significantly different than zero, t(198) = −10.9, p < .001. A chi-square goodness-of-fit test further confirmed that the distribution of errors was significantly different than chance (equal number of responses above and below zero), 𝜒^2^ (1, 𝑁 = 195) = 85.3, 𝑝 < .001: the majority of errors were negative.

Figure 4b shows error as a function of hue distance from start. There was a negative correlation between hue distance and error (error = 0.024 -0.647*distance): as the hue distance from start increased, error became more negative. A linear regression model reveals this trend is significant, r(197) = .448, t = −7.04, p < .001.

46 out of 199 (23.1%) observers reported noticing the change. Thus, the vast majority of observers experienced change blindness and also reported serially dependent hue judgments. Figure 4 (c & d) replots the data separated by noticed response. On the top right of Figure 4c is a plot of two histograms overlayed (yes or no noticed) representing the distribution of errors. A linear regression fit to the yes-noticed data (selected = 288.2 + 0.077*distance) did not find hue distance to be a significant predictor of selected hue, r(44) = .045, t = 0.319 p = .751. A linear regression fit to the no-noticed data (selected = 278.0 + 0.415*distance) did find hue distance to be a significant predictor of selected hue, r(151) = .326, t = 4.24, p < .001 (Figure 4c). A Kolmogorov-Smirnov test indicated that the overall distributions of distance in the yes-noticed (N=46) and no-noticed (N=153) were not significantly different, D(46,153) = .174, p = .204 (Figure 4e). This indicates that longer exposure duration did not increase the likelihood of reporting the change.

## Discussion

In Experiment 2, the slope of selected hue as a function of morph distance was significantly different than zero, and significantly different than ground truth (1). This replicated the main findings from Experiment 1: observers updated their representations over time, but these representations were biased towards the past. The slope of the regression line was 0.353, indicating that observers’ representation of the actual hue at any moment was 35.3% of the true hue at that moment. The distribution of errors had a negative mean error and was not symmetric around zero, which further highlights the bias in observers’ judgments towards earlier hues. Experiment 2 also provided additional evidence against hysteresis and implicit updating, showing that the error became more negative as the amount of preceding information increased (Figure 4b). Given the updated task asking observers to specifically make a report based on the final couch hue, we are confident that our results are not due to averaging. Together, these findings support the idea that observers’ percepts at each moment of a slowly changing stimulus are serially dependent.

Asking observers if they noticed the change provided confirmation that a large majority (76.9%) of the observers who reported these biased percepts (exhibiting evidence of serial dependence) were indeed experiencing slow change blindness. This strengthens the interpretation that continuously serially dependent percepts can produce slow change blindness. To further qualify this interpretation, the data from yes-noticed and no-noticed groups were analyzed separately. The pattern of overall results holds when just considering the no-noticed group: when evaluating how well distance predicted selected hue for the no-noticed data separately, distance was a significant predictor. For the yes-noticed group, the linear regression did not yield a significant relationship between distance and selected hue. However, given the low number of yes-noticed observations, there is not enough evidence to draw concrete conclusions about the relationship between distance and selected hue among the yes-noticed observers, nor can the yes-noticed and no-noticed data be directly compared. Although distance predicts error across all observers, the data collected is not sufficient to conclude that noticing the change results in a difference in reported error. Nonetheless, observers’ hue reports are biased towards the past regardless of whether they report noticing the change.

Intuitively, one might expect that more serial dependence (larger error) leads to slow change blindness while less serial dependence (smaller error) would lead to noticing the change. However, it is unclear where in the experiment time course the yes-noticed observers perceived the change. For example, an observer may notice a change in hue at, say, 10 seconds into the morph. The morph may proceed for another 30 seconds - during which the observer now begins to experience serially dependent percepts, masking any future changes. Now, the observer reports their percept at 40s, which is biased towards the past, and reports that they noticed a change during the trial. In another case, an observer may generate a biased percept that no longer matches the actual hue yet may be sufficiently different from the starting hue to prompt noticing a difference. Their reported percept is serially dependent, yet they have overcome the blindness to the change. From the data collected, it is not feasible to distinguish between the two situations, preventing a comparison between the yes-noticed and no-noticed groups directly. However, the finding that no-noticed observers have serially dependent representations is compelling evidence that slow change blindness is produced from serial dependence.

### Relation to Experiment 1

Both Experiments 1 and 2 provide evidence that perception at any moment while viewing a slow change is serially dependent: reports of stimulus hue are heavily contaminated by the recent past. Error in reported hue increases as the morph proceeds, ruling out hysteresis and implicit updating. This effect holds across two distinct hue ranges, together resulting in nearly 400 unique participants who indicate robust evidence of the effect.

Though the overall results of Experiment 1 and Experiment 2 agree, the slopes differed slightly between the two experiments. There were a number of differences between the two experiments that preclude a meaningful comparison. The pairs of hues used in the two experiments span the same “distance” in HSV hue space but may not have the same linearity or perceptual distance. This is extremely difficult to control for in online experiments, which was known when designing the experiment. An online experiment yielding a large amount of one-shot data at the expense of control of color was a better trade-off than highly controlling color and attempting to recruit hundreds of in-person participants for a minute-long one-shot trial.

The wording of the final question also differed between the two experiments. In Experiment 1, the couch was removed from the screen before the question was asked but the final question appeared while the couch was on screen in Experiment 2. The purpose of having the question appear before the couch was removed was to ensure participants were choosing which hue matched the ending couch, rather than reporting an overall average or simply the initial couch hue. Regardless of these design differences, we draw the same conclusions from both Experiment 1 and Experiment 2.

## Experiment 3

If serial dependence causes the biases in response hue seen in Experiment 1 and Experiment 2, then reducing serial dependence should reduce the biases in response hues. A well-known property of serial dependence is that it is object-selective: when an object changes identity, serial dependence is reduced (Collins, 2020; Liberman et al., 2014; Pascucci et al., 2023; Valdez et al., 2018). Response bias and anchoring, on the other hand, are not object selective. In Experiment 3, observers viewed a slowly changing couch but were then asked to report the color of a different object – a boat. The boat was identical in hue to the ending couch used in Experiment 2, allowing a direct comparison of participant responses between the two experiments. Therefore, if serial dependence drives the observer’s response, hue reports to a *different* object following the same slow change should show less serial dependence (smaller overall error, steeper selected hue slope). If the pattern observed in Experiment 2 is the result of something other than serial dependence, such as response bias or anchoring, there should be no difference in the selected hue slopes between Experiments 2 and 3.

## Method

### Participants

198 new participants were recruited through Prolific (Palan & Schitter, 2018; www.prolific.com) and completed the study online through Pavlovia (Peirce et al., 2019; www.pavlovia.com) for pay at the rate of $10/hr. Participants were required to use a desktop computer but there was no other exclusion criteria. The study was approved by the University of California, Berkeley IRB and informed consent was given by each participant before beginning the study.

### Procedure

The procedure was identical to Experiment 2 in all other ways than described below. The instructions in Experiment 3 were altered slightly to better align with the updated test question. The new instructions read: “You’ll see a colored image of a scene for a random duration, up to 40 seconds. Please pay attention to the image. At some point a black outline will surround the image and you will be asked a question about the scene. Try to be as accurate as possible!

When you are ready, press ‘space’.” Each participant completed only a single trial. The trial began with the slowly changing “test couch” (Figure 5a). At a pre-determined random moment, the couch image was changed immediately to an image of a boat (“test boat”) where the boat hue maintained the planned couch hue. For the last second of the morph (1.5 degrees of hue change) and two additional seconds of static hue, observers viewed the boat instead of the couch. A black outline box and the words “What is the color of the boat?” appeared on the screen for these three seconds. The boat and question then disappeared, and the color wheel and text with a “response boat” were presented to collect the color judgement. Participants used the color wheel to report on the color of the boat rather than the color of the couch. After the participant submitted their color choice, they were then asked: “While viewing the couch on the screen, did you notice any change in the color of the couch?” and responded by clicking a “yes” box or a “no” box on the screen. Participants were then asked about the confidence of their answer, choosing “Total guess”, “Not very confident”, “A little confident”, or “Very confident” (coded as 1,2,3,4). Morph duration (time), morph distance (hue), actual ending hue, selected ending hue, whether the change was noticed, and observers’ confidence were recorded for each participant’s trial.

**Figure 5.**
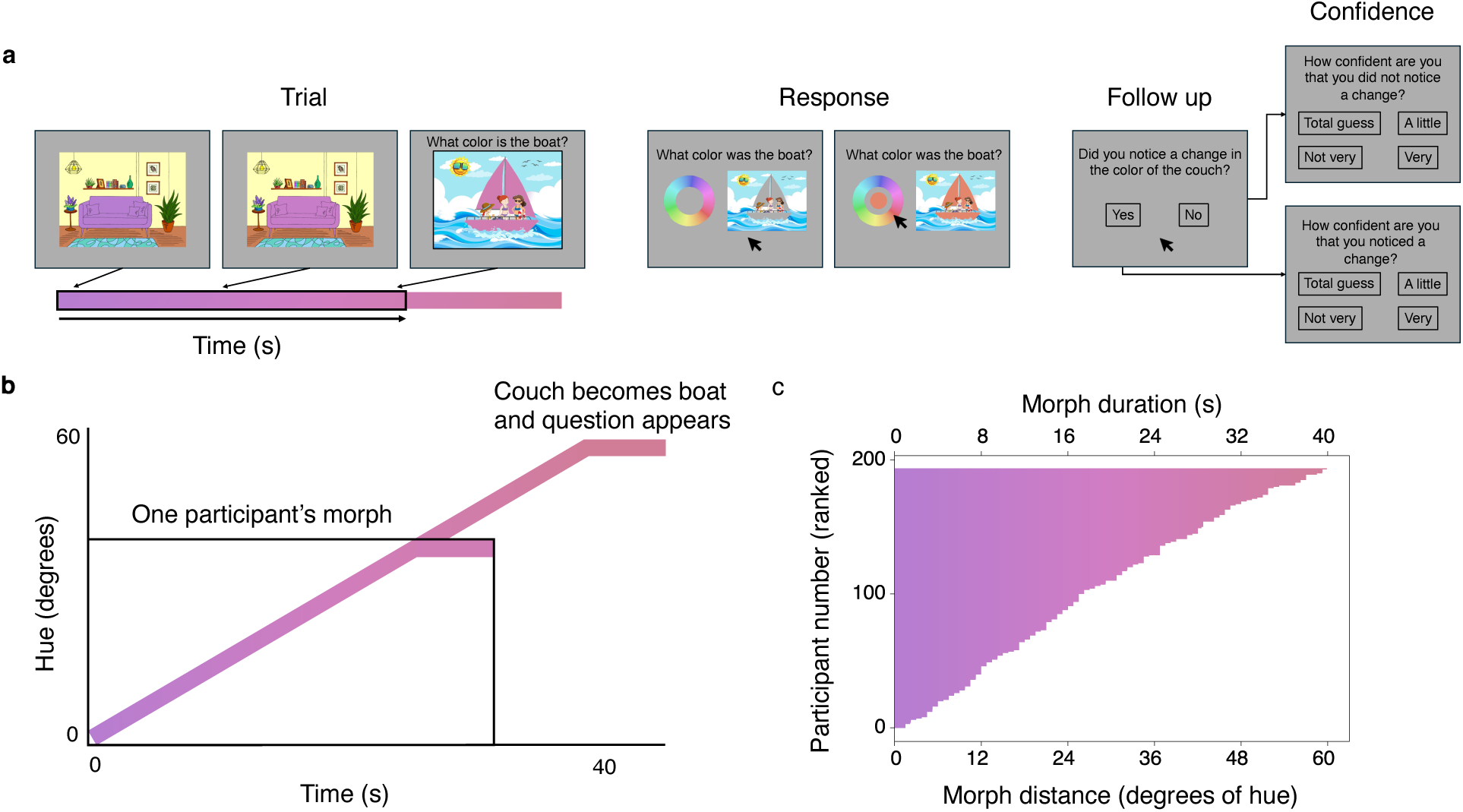
**Experiment 3 stimulus & design**. **a.** The trial consisted of a test couch that changes hue across time. On the left, the test couch is shown in its initial state. The color gradient below the couch represents the maximum hue change possible, with the black outline showing the random subset of the morph that an example participant would experience. The arrow points to the current hue of the couch in each frame. The trial frame on the right shows that the couch has suddenly become a boat, and participants are asked about the color of the boat. On the right is the response screen. Participants used their mouse to hover over the color wheel. As they hovered over the wheel (right), the center of the color wheel and the response boat match the color under the mouse. Participants clicked with their mouse to select their choice. They were then asked whether they noticed a change, and to report their confidence in the decision. **b.** A plot showing stimulus motion (hue change) across time, with an example participant’s morph outlined in black. **c.** The extent of the morph experienced by each participant. The bottom x-axis shows the duration of the participant’s morph, with the corresponding hue distance on the top x-axis. The y-axis shows individual participants. A single row represents what a single observer saw, with the rightmost edge of the bar indicating the actual ending hue at the moment the observer was asked to make a response.

The couch changed at a rate of 1.5 degrees/second and spanned the same range of hues as Experiment 2 (always beginning at 280, travelling a maximum of 60 degrees; Figure 5b). Each individual participant was exposed to a different and random degree of hue change during the single trial (Figure 5c).

### Preprocessing

For each participant, error was calculated as reported hue – actual hue. Raw errors spanned the range −220 to 104. Participants with an error greater than three times the standard deviation above the mean or less than three times the standard deviation below the mean were excluded from analysis and are not depicted in any figures (4 participants).

Before beginning the task, participants were asked to report whether they had normal or corrected-to-normal (wearing glasses or contacts) vision, and whether they were colorblind. From these responses, participants were split into an “impaired” (N = 56) and a “sighted” (N = 138) group. A permutation analysis was conducted to determine whether the difference in mean error between the “impaired” and “sighted” groups was significant. The empirical mean difference (−2.46) was compared to a permuted null distribution, testing the null hypothesis that there is no difference between the means. To create the null distribution, the experiment labels were randomly permuted 10,000 times and the slope was recalculated at each permutation.

The resulting p-value was .683 (α = 0.0167), indicating that the empirical mean difference was not significantly different than zero. Therefore, no data was excluded based on sight during the following analyses.

## Results

Figure 6a shows selected hue as a function of relative hue distance from start. The solid black line reflects a linear regression fit to the empirical data (selected = 280.2 + 0.793*distance) which found distance to be a significant predictor of selected hue, r(192) = .355, t = 5.25, p < .001. An interaction analysis was performed to determine whether the empirical slope (solid black) differed from the unity line (dotted black). A multiple linear regression model (selected ∼ distance*group) fit to the combination of both the ground truth (group 0) and empirical (group 1) data did not yield a significant interaction term (distance:group), meaning that the effect of distance on selected hue did not differ depending on the group, t = −1.37, p = 0.171. This indicates that the slope of the selected hues does not differ from ground truth. The model also yielded a significant main effect of distance (t = 9.37, p < .001) and a non-significant main effect of group (t = 0.032, p = .975).

**Figure 6.**
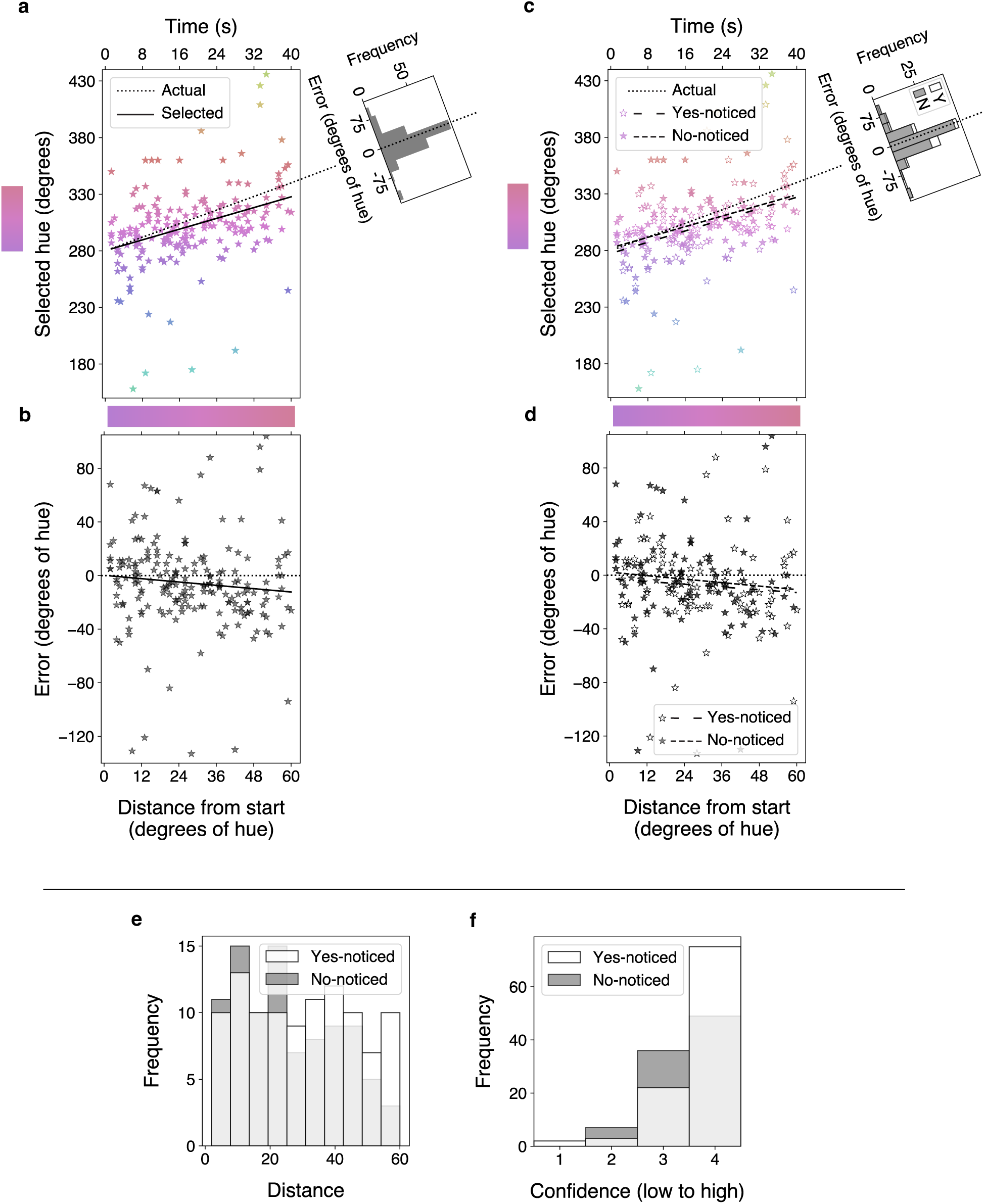
Experiment 3 results. **a.** Selected hue is plotted as a function of the hue distance travelled. Equivalent time is shown on the top x-axis. One star is one participant. The x-axis position reflects the relative morph endpoint, and the horizontal gradient bar shows the actual color at each hue distance. The y-axis position, also the color of the star, is the selected hue. The dotted black line is a unity line representing ground truth: the actual hue at each distance. Any star that does not fall on the dotted line indicates a nonzero error. The solid black line is a regression line fit to the selected hue data. The histogram in the top right corner plots the distribution of errors (selected hue – actual hue). The extended unity line represents zero error. **b.** Error is plotted as a function of hue distance from start, with equivalent time on the top x-axis. Each star represents one participant. The dotted black line is ground truth and the solid black line is a linear regression fit to the error data. **c & d.** Replicate panels a & b with data split by yes-noticed (open; long dash) and no-noticed (filled; short dash). **e.** Frequency of morph distance split by noticed response (yes or no). **f.** Frequency of confidence ratings split by noticed response. A histogram of errors collapsed across the morph is presented in the top right of Figure 6a. A one-sample t test found that the mean error (−5.57, SD = 34.1) is significantly different than zero, t(193) = −2.27, p = .024. A chi-square goodness-of-fit test revealed that the distribution of errors was significantly different than chance, 𝜒^2^ (1, 𝑁 = 188) = 10.3, 𝑝 = .001, indicating that observers overall report an earlier stimulus state.

Figure 6b shows error as a function of hue distance from start. A linear regression model (error = 0.153 −0.207*distance) indicates that this trend is not significant, r(192) = .099, t = −1.37, p = .171: there is no significant relationship between reporting error and hue distance travelled.

102 out of 194 (52.6%) observers reported noticing the change. Figure 6 (c & d) replots the data separated by noticed response. On the top right is a plot of two histograms overlayed (yes or no noticed) representing the distribution of errors (Figure 6c). A linear regression fit to the yes-noticed data (selected = 277.4 + 0.827*distance) found hue distance to be a significant predictor of selected hue, r(100) = .387, t = 4.20 p < .001. A linear regression fit to the no-noticed data (selected = 282.3 + 0.783*distance) also found distance to be a significant predictor of selected hue, r(90) = .326, t = 3.27, p = .002 (Figure 6c). A Kolmogorov-Smirnov test indicated that the overall distributions of distance in the yes-noticed (N=102) and no-noticed (N=92) were not significantly different, D(102,92) = .148, p = .212 (Figure 6e), indicating that a greater exposure to the morph did not increase the likelihood of reporting the change. A chi-square test for independence revealed that there were significantly more observers who reported noticing the change in Experiment 3 compared to Experiment 2, 𝜒^2^ (1, 𝑁 = 393) = 35.1, 𝑝 < .001.

The majority of observers reported that they were highly confident, both those who reported noticing the change and those who reported not noticing it (Figure 6f). A Kolmogorov-Smirnov test indicated that the overall distributions of confidence responses in the yes-noticed (N=102) and no-noticed (N=92) were significantly different, D(102,92) = .203, p = .031.

Figure 7a replots the data from Experiment 2 (diamond, black) and 3 on the same plot (star, gray), which differed only in the identity of the object at the end of the morph and during report. On the top right is a plot of two histograms overlayed (from each experiment) representing the distribution of errors. A permutation analysis was conducted to determine whether the difference in slope of reported hue across the morph between Experiment 2 (slope = 0.353) and Experiment 3 (slope = 0.793) was significant. The empirical slope difference (0.440) was compared to a permuted null distribution, testing the null hypothesis that there is no difference between the slopes. To create the null distribution, the experiment labels were randomly permuted 10,000 times and the slope was recalculated at each permutation. The resulting p-value was .010 (α = 0.05), indicating that the empirical slope difference was significantly different than chance (Figure 7b). A Kolmogorov-Smirnov test found that the overall distributions of error in Experiment 2 (N=199) and Experiment 3 (N=194) were significantly different, D(199,194) = .216, p < .001, indicating that error in Experiment 3 was overall smaller and more centered around zero. Together, this indicates that the change in object identity introduced in Experiment 3 significantly reduced serial dependence.

**Figure 7.**
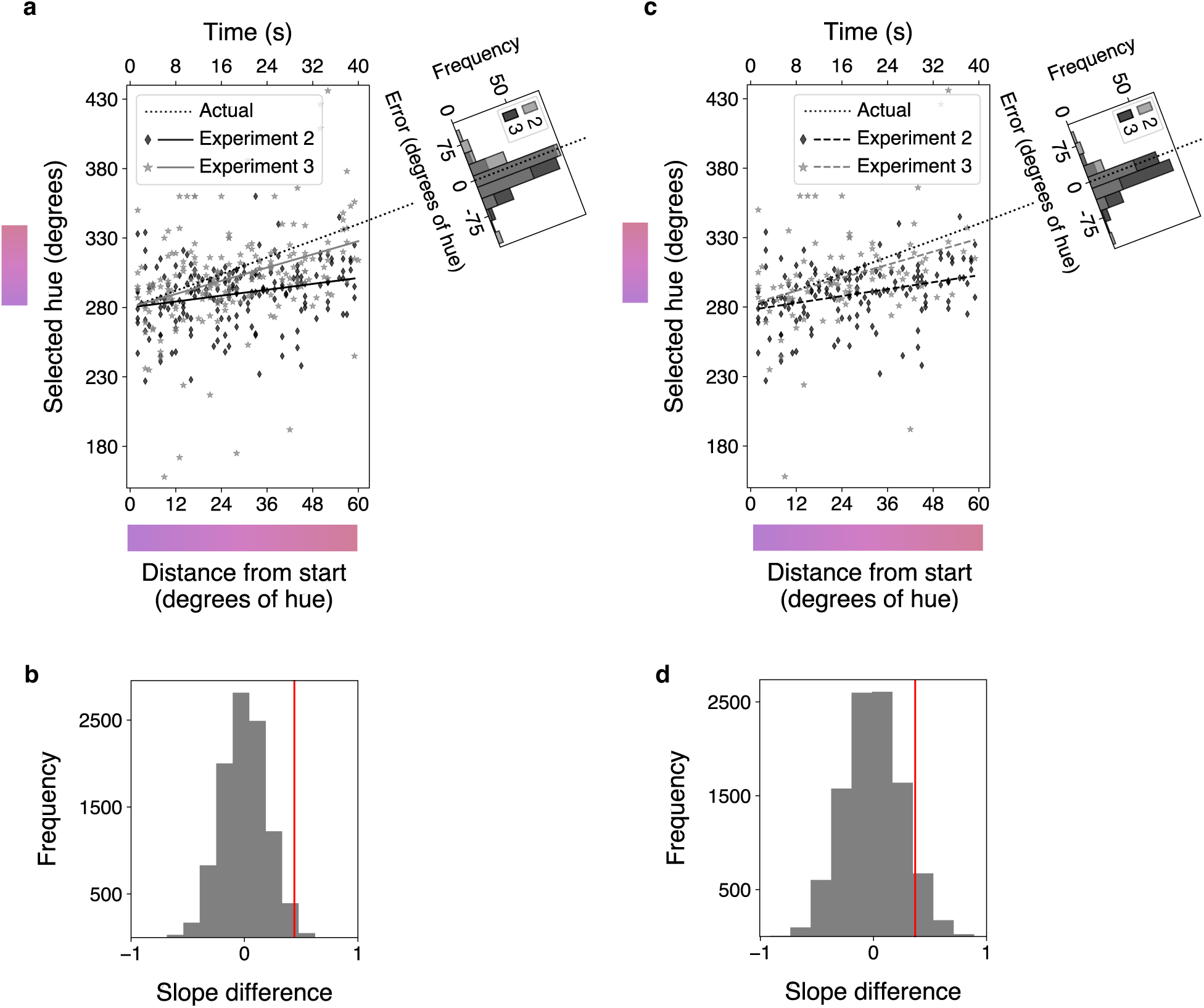
The results from Experiments 2 (diamond, black) and 3 (star, gray) on the same plot (combining Figures 4 and 6). **a.** All observers from the two experiments. **b.** Permuted null distribution of slope difference between all observers in Experiments 2 and 3. Red line is empirical slope difference. **c.** Only the no-noticed observers from Experiments 2 and 3. **d.** Permuted null distribution of slope difference between only the no-noticed observer in Experiments 2 and 3. Red line is empirical slope difference.

Figure 7c replots only the no-noticed data from Experiment 2 (diamond, black) and 3 on the same plot (star, gray). A permutation analysis was conducted to determine whether among the no-noticers the difference in slope across the morph between Experiments 2 (0.415) and 3 (0.783) was significant. The empirical slope difference (0.368) was compared to a permuted null distribution, testing the null hypothesis that there is no difference between the slopes. As above, the experiment labels were randomly permuted 10,000 times and the slope difference was recalculated. The resulting p-value was .075 (α = 0.05), indicating that there is not enough evidence to reject the null hypothesis (Figure 7d). There is a strong trend in the data to support that the slopes are different, but there is not enough power due to the lower proportion of no-noticed trials in Experiment 3. A Kolmogorov-Smirnov test found that the overall distributions of error among non-noticers in Experiment 2 (N=153) and Experiment 3 (N=92) were significantly different, D(153,92) = .282 p < .001, indicating that even within the non-noticers, error in Experiment 3 was overall smaller and more centered around zero.

## Discussion

In Experiment 3, the slope of selected hue as a function of morph distance was significantly different than zero, but not significantly different than 1 – the ground truth unity line. In other words, observers accurately reported the color of the boat. Though the mean error and distribution of errors were significant, there is no effect of distance on hue error. The main goal was to compare Experiment 3 to Experiment 2, which were identical save for the object identity change. The error distribution in Experiment 3 was significantly different than in Experiment 2. This suggests that the smaller mean error in Experiment 3 is the result of the distribution becoming more centered around zero, evidence of reduced bias in selected hue reports. Further, selected hue across the morph had a significantly steeper slope in Experiment 3 compared to Experiment 2. The reduction in bias in Experiment 3 confirmed that observers’ responses in Experiment 2 were object-specific, rather than a hue in general, and therefore driven by serial dependence. We conclude that active perceptual serial dependence integrates similar information across time to maintain a stable percept of the couch and that when the couch suddenly becomes a boat, the information is too different to be reconciled: the boat, and its current hue, become the new stable reference point.

Participants in Experiment 3 were also asked whether they noticed the change. Surprisingly, 52.6% of observers reported yes - a significant increase from Experiment 2. Statistical analysis found there was no effect of observers’ reported noticing on the pattern of errors, consistent with the findings from Experiment 2. However, despite the question asking specifically about a change in the couch, it is possible that the change which participants reported noticing was actually the change from the couch to the boat. While this is purely speculative, this could explain the near doubling of reported noticers. If this speculation is true, noticing a change between the couch hue and boat hue when the true difference is less than 1.5 degrees could further strengthen the interpretation that observers have a percept of the couch that matches the past. The sudden appearance of the boat may introduce a temporal or event boundary prompting a re-assessment of the features of the object, which are no longer serially dependent. If the percept of the couch is biased and the percept of the boat is not, it stands to reason that despite being nearly identical, observers perceive a large change in the hue between the two objects.

Given the large proportion of yes-noticed observers in Experiment 3, an additional comparison was made between only the no-noticed observers in Experiments 2 and 3 (Figures 10b and 10d). The overall error histograms were significantly different, demonstrating the overall increased accuracy among no-noticers in Experiment 3. A visible comparison of the selected hue slopes (Figure 10b) shows a striking difference, yet a permutation analysis did not yield enough evidence to conclude that the two slopes were significantly different. The increased variance in the no-noticed data from Experiment 3, likely due to the lower proportion of data included, may explain the lack of power. Together with the differences in the overall data between Experiments 2 and 3, the trending difference between the no-noticers does support the conclusion that the bias in responses is serial dependence.

## Experiment 4

Though the choice to use one-shot responses was intentional and useful, we remained curious whether observers who knew about the task and the nature of the slow changes would still generate serially dependent percepts. We therefore conducted a version of Experiment 1 in the lab. This would allow us to evaluate whether the effect could be replicated with repeated trials and address any concerns about the effects of the specific hues used in Experiments 1-3. If we did find a pattern of serial dependence in Experiment 4, we hypothesized that the effect size would be much smaller as a result of participants knowing about the task and stimulus. If we did not find the pattern in Experiment 4, this would not negate the results of Experiments 1-3 but it might suggest that observers change their integration strategy when information is known to change.

## Method

### Participants

12 observers were recruited through the University of California, Berkeley’s Psychology Department Research Participation Program and completed the study for credit (1 credit/hour). The study was approved by the University of California, Berkeley IRB and all participants provided written informed consent prior to participation in the study.

### Procedure

The procedure for this experiment closely followed the procedure from Experiment 1, but a few changes were made to account for the repeated trials. The 12 observers completed an average of 195 trials each (Figure 8). 16.67% of the trials included no change. This was to introduce uncertainty and ensure that participants were paying attention throughout the morph and were not included in analysis. Colors were defined in HCL space with a constant chromaticity (35) and lightness (70). The starting hue was chosen randomly and the distance varied randomly on each trial and images were presented on a gamma corrected and color calibrated monitor. The hue changed at a rate of 2 degrees per second for all trials (this is faster than the online studies and was based the results of a separate in-lab task). Each trial began with the image of the couch on the screen. The couch remained on the screen for a random duration and may or may not have changed hue. After the couch was removed from the screen, participants were asked to use their mouse to select the hue that best matched the couch they had just seen. The orientation of the color wheel was randomized on each trial. Once the participant submitted their response, there was a brief 100 ms delay and the next trial began.

**Figure 8.**
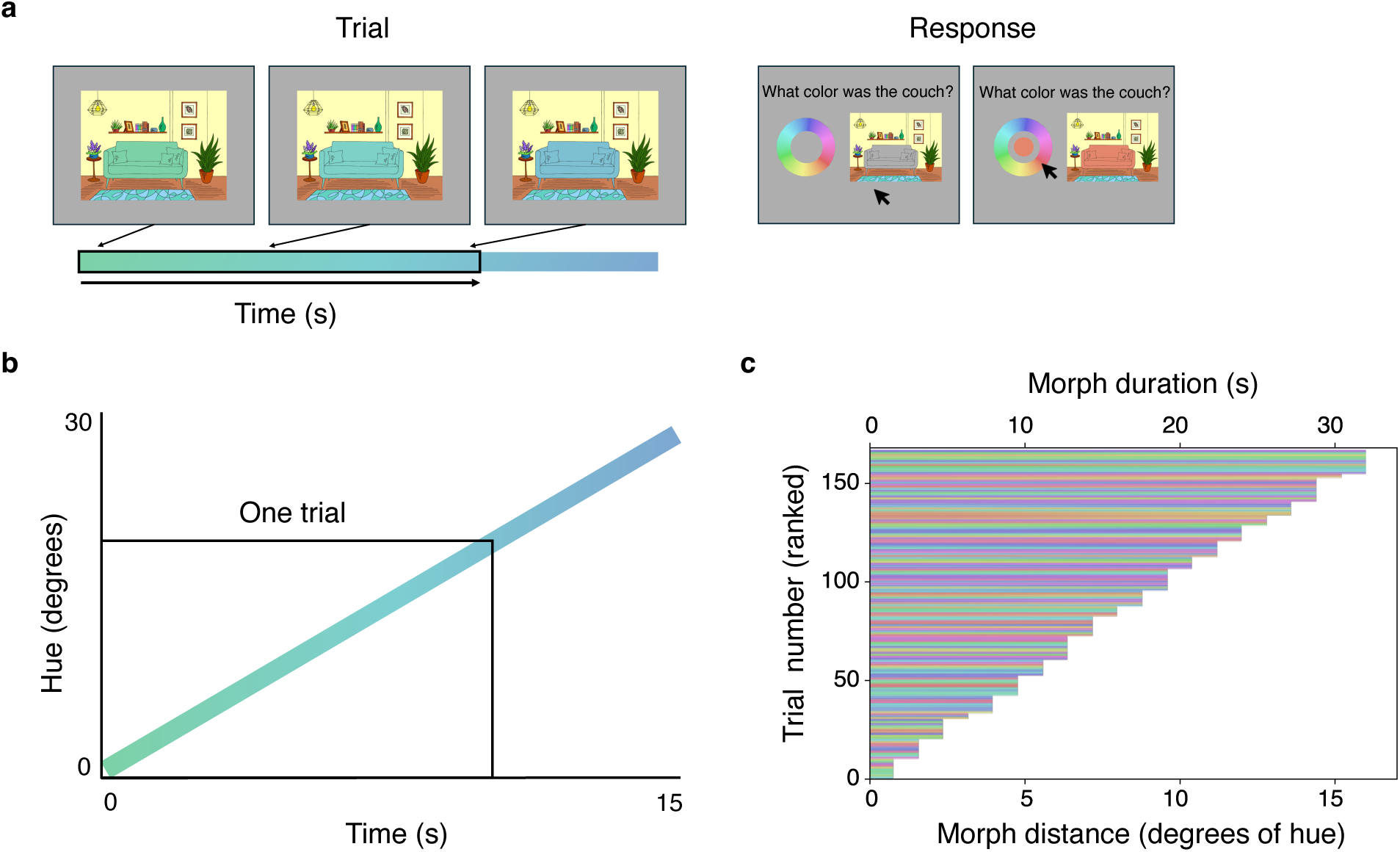
**Experiment 4 stimulus & design**. **a.** A schematic of a trial, which was identical to Experiment 1. **b.** A plot showing stimulus motion (hue change) across time, with an example trial outlined in black. **c.** Duration of morph on each trial experienced by an example participant. All trials for an example participant are plotted, ranked by distance. The varying colors of the horizontal bars reflect the actual hues used in each trial; starting hue was chosen randomly on each trial.

The task was the same for each trial. As such, participants knew from trial 2 onward that they would explicitly be asked about the color of the couch. Further, the participants knew that the couch color could change. Aside from the slight stimulus modifications and the nature of the multi-trial design, the in-lab experiment was identical to Experiment 1. For analysis, the starting hue on each trial was standardized to be 0, and actual ending hue and reported ending hue were redefined with respect to this new baseline. Reported ending hues were adjusted to fall within +/-180 degrees of the starting hue (for example, a reported hue of 200 is equivalent to a reported hue of −160). Error was calculated as reported hue – actual hue. Trials with an error greater than three times the standard deviation above the mean or less than three times the standard deviation below the mean were excluded from analysis and are not depicted in any figures (36 trials).

## Results

Figure 9a shows relative selected hue (trials standardized so that starting hue equaled 0) as a function of hue distance from start. A linear regression fit to the in-lab data (selected = −0.596 + 0.817*distance), the solid black line, found distance to be a significant predictor of relative selected hue, r(1922) = .432, t = 21.0, p < .001 and an interaction analysis (selected ∼ distance*group) revealed a significant difference between the empirical slope (solid black) and the unity line (dotted black), t = −4.71, p < .001 (Figure 13a). This means that the slope of the selected hues was significantly flatter than ground truth. The multiple regression model also yielded a significant main effect of distance (t =36.4, p < .001) and a non-significant main effect of group (t = −0.923, p = .356).

**Figure 9.**
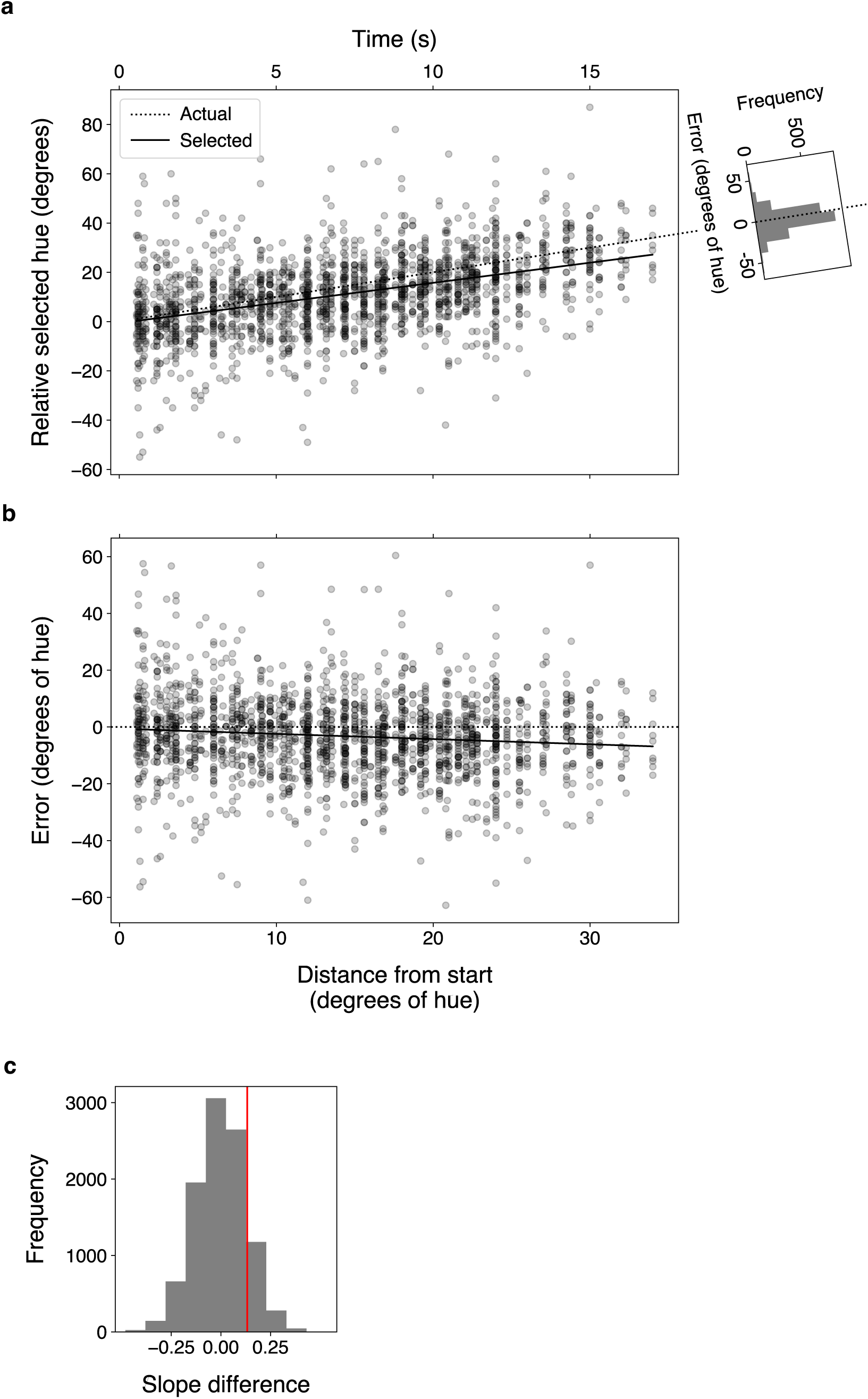
Experiment 4 results. **a.** Relative selected hue (all trials standardized to have a starting hue of zero) plotted as a function of hue distance from start; equivalent time plotted on top x-axis. One circle is one trial. Data from 12 participants is shown here. The x-axis position reflects the relative morph endpoint, and the y-axis position is the selected hue relative to a starting hue of zero. The dotted black line is a unity line representing ground truth: the actual hue relative to start at each distance. Any circle that does not fall on the dotted line indicates a nonzero error. The solid black line is a regression line fit to the selected hue data. The histogram in the top right corner plots the distribution of errors (selected hue – actual hue). The extended unity line represents zero error. **b.** Error is plotted as a function of hue distance from start, with equivalent time on the top x-axis. Each circle represents one trial; data from 12 participants is shown. The dotted black line is ground truth and the solid black line is a linear regression fit to the error data. **c.** Permuted null distribution of slope difference between Experiments 1 and 4. Red line is empirical slope difference.

The overall mean error was negative (−3.24, SD = 14.1), which a one-sample t test found was significantly different than zero, t(1923) = −10.0, p < .001 (histogram to the right of Figure 9a). A chi-square goodness-of-fit test revealed that the distribution of errors was significantly different than chance, 𝜒^2^ (1, 𝑁 = 1899) = 97.8, 𝑝 < .001, indicating that observers report an earlier stimulus on a significant majority of trials. The slope of the relationship between error and hue distance (−0.183) was significant, r(1922) = .105, t = −4.71, p < .001 (Figure 9b).

The in-lab effect size seemed to be more modest than we found online. A permutation analysis was conducted to determine whether the slope difference between Experiment 1 (0.684) and Experiment 4 (0.817) was significantly different. The empirical slope difference (0.133) was compared to a permuted null distribution, testing the null hypothesis that there is no difference between the slopes. The experiment labels were randomly permuted 10,000 times and the slope difference was recalculated. The resulting p-value was .142 (α = 0.05), indicating that there is not enough evidence to reject the null hypothesis (Figure 9c). Together, this hints that knowledge about a stimulus’ potential to change does not prevent active perceptual serial dependence from biasing perception.

## General Discussion

We used single-observer one-shot trials to evaluate biases in perception of slowly changing objects across time. Consistent with previous work (Frey, Koenig, Block, et al., 2024; Laloyaux et al., 2008; Manassi & Whitney, 2022), after viewing an object that slowly changes hue, observers report that object’s hue to match an earlier state rather than the true end state (Experiments 1, 2, and 4). Expanding on previous work, we further reveal that these perceptual biases exist at each moment throughout the slow change, and that the magnitude of the bias depends on the preceding content – becoming greater as the current state evolves farther from the initial one (Experiments 1, 2, and 4). In Experiment 3, there is a significant reduction in serially biased hue reports when the object changes identity at the end of the morph: confirmation that the bias in hue reports is in fact serial dependence rather than anchoring, hysteresis, or response bias. Together, these results provide evidence in favor of the active perceptual serial dependence hypothesis (Manassi & Whitney, 2022).

### Continuity fields

These results support the idea that an on-line stability mechanism, active perceptual serial dependence, smooths perception: the more of the morph observers saw, the larger the bias towards the past. For participants who viewed the longest duration of morph (40s), their reports most closely match the stimulus as it was around 13s ago (Experiment 1) and at around 27s ago (Experiment 2). This suggests that information presented roughly 20 seconds ago can shape current perception, consistent with estimates of continuity fields in previous research (Manassi et al., 2023). However, based on our results alone, we are unable to determine whether the continuity field is determined by time, difference in stimulus features, the rate of change, or a combination of these factors. Further, serial dependence induced slow change blindness may not be complete. For most observers, the smoothing was sufficient to produce slow change blindness. However, successive information that exceeds the bounds of the continuity field may cause observers to notice a change. If the target feature changes too quickly, each successive stimulus state will be sufficiently different to override a stabilization mechanism. When changes occur more quickly, they also may introduce a visual transient that triggers detection. Likewise, an object identity change like in Experiment 3 may serve as an event boundary (Radvansky & Zacks, 2017; Swallow et al., 2009) which triggers detection.

Observers might also switch from internal to external modes as they accumulate increasingly different information, prompting them to detect the change (Weilnhammer et al., 2024). Future work should consider evaluating the pattern of error responses across time to stimuli that change at different rates.

Why might the visual system strive for continuity at the expense of detecting changes? It is computationally efficient and capitalizes on the overall correlation of natural world statistics over time (Dong & Atick, 1995). It is unlikely that a couch will start slowly changing color.

Therefore, using recent perception of the couch color to inform current perception often yields an accurate representation. In other cases, any variation in the color that is not noticed is likely to be due to changes in the surrounding illumination (Foster, 2025). To interpret these variations as changes in the couch’s feature would be costly, slow, and inefficient. So, is a better strategy to assume stability and miss gradual changes, or assume everything is changing and struggle to adapt to such changes? Our visual system seems to choose the former, at least in some circumstances.

### Value of one-shot experiments

The approach used in our experiments relied on single-observer reports to different morph durations, which, stitched together, sampled an entire morph range. While each observer was completely independent, together their pattern of responses yields a proxy for perception across time. The one-shot approach ensured that participants viewed the slowly changing couch with no knowledge of the end task, no knowledge that the couch could change, and no knowledge of the range of hues the couch traversed. Therefore, there was no contamination of task-goals or expectation, and observers behaved naturally during viewing and response. Repeated trials would risk spoiling the slow change, alter participant strategy during morph viewing, and prevent us from answering our desired question. Nevertheless, Experiment 4 did confirm that the general pattern of results holds even when observers do have prior knowledge of task, design, and stimulus.

### Importance of attention

It is widely agreed upon in the classic change blindness field that attention is necessary for change detection (Hollingworth & Henderson, 2002; Rensink et al., 1997). An explanation of the high rate of slow change blindness might simply be that observers were not paying attention to the stimulus, and instead experienced inattentional blindness (Simons & Chabris, 1999). For example, observers who have been watching a seemingly unchanging couch might stop paying attention at some point, and when asked to report the color of the couch, have only a representation from when they were attending to report on. However, the high precision of hue judgments and high confidence ratings suggest that the participants did attend to the changing stimulus. Participants knew they needed to continuously attend to the stimulus because they would be asked a question about the stimulus when the trial ended, which would happen randomly but not exceed 40 seconds in length. Further, observers who do not attend to the changing object should not maintain a representation of it. Therefore, when prompted to report on the final stimulus state in Experiment 2, their attention would be drawn to the current ending state and their report would have little bias. This suggests that inattention would predict a result similar to the implicit updating hypothesis, which is not what was found here. Overall, our results suggest that observers did pay attention.

How is it possible that our observers were attentive, yet still failed to notice the change? Some research has found that attention is not always enough: observers fail to notice changes to the object they are attending and even fixating (Levin & Simons, 1997; Simons & Levin, 1998). Further, serial dependence depends on attention (Fischer & Whitney, 2014; Fu & Mei, 2024; Kiyonaga et al., 2017b), with greater attention leading to greater serial dependence (Manassi et al., 2023). Given that our results provide evidence that slow change blindness arises from serial dependence, it stands to reason that our observers are more blind *because* of their attention, not because of a lack of it. As observers attend to the changing scene, serial dependence works to stabilize the impression, biasing perception and camouflaging the change.

### The importance of memory

Throughout this investigation, we have taken observer reports as a proxy for perception at each moment. However, the report task in Experiment 1 likely contained some influence of memory. In Experiments 2 and 3, we updated the task to ask observers to make a judgement about a particular stimulus while it was on the screen. While serial dependencies in perception may be considered a form of memory, the updated task in Experiments 2 and 3 greatly reduced the contributions of longer-term memory stores to participants’ responses. Even if we did probe memory here, not only are these results consistent with previous work probing memory during slow change blindness (Frey, Koenig, Block, et al., 2024), but these results still show robust evidence of serial dependence regardless of whether observer responses include dependencies in memory.

## Conclusion

Integrating information across time is a fundamental task for the visual system. Here, we provide evidence that the visual system combines information from the recent past with current stimulus information to generate perception. This active perceptual serial dependence is an on-line mechanism that assumes, and in turn promotes, visual stability in features that might actually be changing. Importantly, we demonstrate a connection between serial dependence and slow change blindness by confirming that the majority of our observers were blind to the slow change. Therefore, we propose that the continuously biased percepts generated by active perceptual serial dependence camouflage the slow change and can lead to slow change blindness.

## Acknowledgments

The authors would like to thank Veith Weilnhammer for recruiting participants and hosting the study online.

## Author contributions

Conceptualization: HGF, DW Methodology: HGF Investigation: HGF Visualization: HGF, DW Writing—original draft: HGF

Writing—review & editing: HGF, DW

## Data and materials availability

All data, code, and materials used are available upon request.

